# A systematic comparison of fibroblasts derived from postmortem human dura mater versus dermal epithelium for neurodegenerative disease modeling

**DOI:** 10.1101/2021.05.17.444554

**Authors:** Andrea R. Argouarch, Celica G. Cosme, Kristle Garcia, Christian I. Corrales, Alissa L. Nana, Anna M. Karydas, Salvatore Spina, Lea T. Grinberg, Bruce Miller, Hani Goodarzi, William W. Seeley, Aimee W. Kao

## Abstract

Patient-derived cells hold great promise for precision medicine approaches in human health. Fibroblast cells have been a major source of human cells for reprogramming and differentiating into specific cell types for disease modeling. Such cells can be isolated at various stages during life (presymptomatic, symptomatic, and postmortem) and thus can potentially be used to model different phases of disease progression. In certain circumstances, however, tissues are not collected during life and only postmortem tissues are the only available source of fibroblasts. Fibroblasts cultured from postmortem human dura mater of individuals with neurodegenerative diseases have been suggested as a primary source of cells for *in vitro* modeling of neurodegenerative diseases. Although fibroblast-like cells from human and mouse dura mater have been previously described, their utility for reprogramming and direct differentiation protocols requires further characterization. In this study, cells derived from dermal biopsies performed in living subjects were compared to cells derived from postmortem dura mater. In two instances, we have isolated and compared dermal and dural cell lines from the same subject. Notably, striking differences between the dermis and dura mater-derived cell lines were found. Compared to dermal fibroblasts, postmortem dura mater-derived cells demonstrated different morphology, exhibited slower growth rates, failed to express fibroblast protein markers, and exhibited significant differences in gene expression profiles. In addition, dura mater-derived cells were found to exhibit a high rate of chromosomal abnormalities, particularly in the loss of the Y chromosome. Our study highlights potential limitations of postmortem human dura mater-derived cells for disease modeling, argues for rigorous karyotyping prior to reprograming, and brings into question the identity of dura mater-derived cells as belonging to a fibroblast lineage.

## INTRODUCTION

The appeal of human *in vitro* cell models stems from the potential to recapitulate and elucidate the pathophysiology of numerous diseases, including neurodegenerative disorders. Patient-derived cell lines can address cellular and molecular pathways that complement other models. The utility of patient-derived cell models depends upon their ability to reliably reflect the disease process. For example, a fibroblast patient-derived cell model of Huntington’s disease has been reported to clinically recapitulate the disease state better than transformed cellular models [15]. In this regard, isolated fibroblasts from routine dermal biopsies are collected in order to be directly differentiated into induced neurons (iNeurons) or reprogrammed into induced pluripotent stem cells (iPSCs) [6,17] that can subsequently be differentiated into neural progenitor cells or different neuronal subtypes. Further incorporation of induced neurons into more complex cellular models, such as three-dimensional brain organoids, represents additional avenues to replicate morphological and functional cell interactions of the human brain [20]. Fibroblasts from dermal biopsies performed in living research subjects have been a common cell source for patient-derived disease models. In certain instances, however, it would also be advantageous to access fibroblasts from postmortem tissue, such as when family members who carry relevant gene mutations are identified or when no other sources are available. For these reasons, an accessible source of postmortem fibroblasts from patients could provide significant value to the field.

Human dura mater can provide an alternative source of primary cells that can be directly obtained from postmortem tissue. Dura mater is the outer most layer of the meninges, located between the skull and the brain, and is primarily composed of fibroblasts and collagen [14]. Cells from dura mater can be isolated, cultured, and utilized for several applications. Previous studies have investigated the use of dura mater-derived cells in grafts [9], drug response [10,12], particle engulfment [18,19], inflammation and healing [5,11], and reprogramming into iPSCs [2,3,22]. Brain biobanks oftentimes collect and store dura mater at the time of autopsy. The use of these dura mater-derived cells can be advantageous because they can be studied when other cell sources, such as dermal fibroblasts and peripheral blood mononuclear cells, were not collected during life. Additionally, dura mater-derived cells as an *in vitro* disease model can provide novel and complementary insights to traditionally established cell-based disease models. Although these dura mater-derived cells have been used in multiple studies, a direct comparison between dermis and dura mater-derived cells has not been reported to our knowledge.

In this study, we endeavored to better characterize the cells isolated from postmortem human dura mater to assess their utility as a source of cells for *in vitro* disease modeling. Sproul and colleagues previously isolated dura mater-derived cells from non-cryoprotected frozen dura mater [22]. Similarly, we isolated cells from postmortem frozen and fresh dura mater from subjects with neurodegenerative diseases and compared cell outgrowth from the tissue, cellular morphology, growth rate, chromosomal karyotype, protein expression of fibroblast markers, and RNA-sequencing profiling to dermis-derived cells. Furthermore, we isolated and characterized dermis and postmortem dura mater-derived cell lines generated from the same subjects. This direct, comparative analysis between cells isolated from the dermis and dura mater highlights the utility and potential limitations of postmortem dura mater-derived cells as an *in vitro* patient-based disease model.

## MATERIALS AND METHODS

### Tissue collection

Dermal tissue was provided by the University of California, San Francisco, Memory and Aging Center, where participants enrolled in observational research studies underwent skin biopsy after providing written informed consent. Approximately two-millimeter dermal punches were performed at the inner thigh. The tissue was immersed in complete media composed of DMEM high glucose with sodium pyruvate (Thermo Fisher Scientific, Waltham, MA, USA, #11995073), 10% heat inactivated FBS (VWR, Radnor, PA, USA, #97068-091), and 1% Penicillin-Streptomycin (Thermo Fisher Scientific, #15140122). Cells were isolated within eight hours after procurement.

Postmortem human dura mater was provided by the University of California, San Francisco, Neurodegenerative Disease Brain Bank. Consent for brain donation was obtained from all subjects or their surrogates, following the principles outline in the Declaration of Helsinki. Prior to brain extraction, the overlying dura mater was removed and stored surrounded by wet ice during transport. Upon arrival to the laboratory, a small patch of dura ranging in maximum dimension from two to five centimeters, was dissected and either (1) immersed in complete media in a centrifuge tube or (2) rapidly frozen and banked. The freshly captured dural tissue was stored at 4°C, and cells were isolated within 24 hours after procurement. For frozen dural tissues, samples were placed in a freezer bag and surrounded by dry ice, then stored in a - 80°C freezer for up to 4 years of long-term without the use of a cryoprotectant reagent. The dural tissue was briefly thawed and cells were isolated.

### Cell culture

Following tissue collection, both dermal and dural tissues were washed in sterile Dulbecco’s phosphate buffered saline (DPBS) without calcium and magnesium (Thermo Fisher Scientific, #14190250) four times, one minute per wash, in a cell culture dish. The tissue was then transferred into a new dish with complete media. The tissue was divided into smaller sections, approximately one to two millimeters, with surgical tools and transferred into a six-well cell culture plate. A glass coverslip was gently placed over the tissue and complete media was added into the wells. For the remaining fresh dural tissue, it was cryopreserved in complete media and 10% DMSO for long-term tissue storage in liquid nitrogen. Cultures were incubated in a humidified chamber at 37°C and 5% CO_2_, left undisturbed for the first week, then fed every two to three days with complete media. After approximately three weeks in culture, the cells were collected, expanded, and banked in liquid nitrogen with complete media and 10% DMSO.

To optimize the recovery of cells from frozen dural tissue, an alternative cell culture media was used, composed of DMEM high glucose with sodium pyruvate, 10% heat inactivated FBS, 1X NEAA (Thermo Fisher Scientific, #11140050), 1X Glutamax (Thermo Fisher Scientific, #35050061), 0.1 mM 2-Mercaptoethanol (Thermo Fisher Scientific, #21985023), 1X Nucleosides (MilliporeSigma, Burlington, MA, USA, #ES-008-D), and 1% Penicillin-Streptomycin [3, 22]. This DMEM media with nucleosides was used for the first week in culture and then switched to complete media with the addition of Glutamax and 2-Mercaptoethanol. Alternatively, DMEM media with nucleosides was used throughout the entire time in culture.

### Immunofluorescence

Cell lines were cultured and plated in a four-well cell culture chamber slide (Thermo Fisher Scientific, #177437). Cell lines included a dermis and frozen dura mater-derived cell line from the same subject and a fresh dural cell line. Upon confluency, cells were washed twice with DPBS with calcium and magnesium (MilliporeSigma, #D8662), fixed with 4% paraformaldehyde (Electron Microscopy Sciences, Hatfield, PA, USA, #15714) for 20 minutes, washed with DPBS^+/+^ 3 times for 5 minutes each, and then permeabilized with 0.1% Triton X-100 (MilliporeSigma, #T8787) for 10 minutes. Cells were washed with DPBS^+/+^ and blocked with 1% bovine serum albumin (Thermo Fisher Scientific, #BP1605-100) and 0.25% Triton X-100 for 30 minutes. Cells were then incubated with 1X rhodamine phalloidin conjugated to TRITC (Thermo Fisher Scientific, #R415) in block solution for one hour and washed with DPBS^+/+^. All solutions were made in DPBS^+/+^ and performed at room temperature. Slides were mounted with Prolong Gold Antifade Reagent with DAPI (Cell Signaling Technologies, Danvers, MA, USA, #8961) as a counterstain and imaged on the DMi8 confocal platform (Leica Microsystems, Wetzlar, Germany) with a 10x objective.

### Cell proliferation

Dermis and dura mater-derived cell lines were plated at the same seeding density in multiple 24-well cell culture plates at day 0. Over the span of six days, cells were counted every day to assess proliferation. Cells were washed with DPBS^-/-^, trypsinized with 0.05% Trypsin-EDTA (Thermo Fisher Scientific, #25300062), and collected for cell counting. All cell counts were determined utilizing the Bio-Rad TC20 automated cell counter (Bio-Rad Laboratories, Hercules, CA, USA, #1450102) with a 1:1 dilution of trypan blue solution (Thermo Fisher Scientific, #15250061) to cell suspension. All cell counts were performed in triplicates.

### Chromosomal karyotyping and mycoplasma analysis

For karyotype analysis, dermis and dura mater-derived cell lines were cultured in a T-25 cell culture flask and shipped live to Cell Line Genetics (Madison, WI, USA) for karyotype analysis using G-banding. For mycoplasma detection, cells were collected, washed with DPBS^-/-^, and DNA was extracted using the QuickExtract DNA Extraction Solution (Epicentre, Madison, WI, USA, #QE09050) according to the manufacturer’s instructions. Mycoplasma PCR detection was performed according to the manufacturer’s instructions (Bulldog Bio, Portsmouth, NH, USA, #25234). All dura mater-derived cell lines were negative for mycoplasma contamination (data not shown).

### Western blot

Dermis and dura mater-derived cell lines were collected, washed with DPBS^-/-^, and lysed in cold RIPA buffer (Thermo Fisher Scientific, #89900) with protease (MilliporeSigma, #4693124001) and phosphatase (MilliporeSigma, #4906837001) inhibitors. Lysates were spun and the supernatants were collected for western blot analysis. Protein concentration was determined by bicinchoninic acid assay according to the manufacturer’s instructions (Thermo Fisher Scientific, #23225). Samples were denatured, separated on a NuPAGE 4-12% Bis-Tris protein gel (Thermo Fisher Scientific, #NP0336BOX), and transferred onto a nitrocellulose membrane. Membranes were blocked in Odyssey blocking buffer (LI-COR, Lincoln, NE, USA, #927-50000) for one hour at room temperature, incubated with primary antibodies overnight at 4°C, and incubated with the appropriate LI-COR secondary antibodies for one hour at room temperature. Membranes were imaged on the LI-COR Odyssey CLx imaging system. The following primary antibodies were used: Rabbit anti-Vimentin (1:1,000, Cell Signaling Technologies, #5741S), Rabbit anti-S100A4 (1:1,000, Cell Signaling Technologies, #13018S), and Mouse anti-Actin (1:5,000, MilliporeSigma, #MAB1501). The following secondary antibodies were used: 680RD Donkey anti-Rabbit (1:10,000, LI-COR, #926-68073), 800CW Donkey anti-Rabbit (1:10,000, LI-COR, #926-32213), and 800CW Donkey anti-Mouse (1:10,000, LI-COR, #926-32212).

### RNA-sequencing

From the same subject, fresh dermal and frozen dural cell lines were plated into a six-well cell culture plate and isolated for RNA-sequencing. Cells were trypsinized, washed with PBS^-/-^, and total RNA was extracted and purified using the Quick-RNA Microprep Kit (Zymo Research, Irvine, CA, USA, #R1051). Samples were extracted in quadruplicates and RNA was quantified using a NanoDrop. Libraries for RNA-sequencing were prepared using the SMARTer Stranded Total RNA-Seq Kit v2 Pico Input Mammalian library preparation protocol (Takara Bio USA, Inc., Mountain View, CA, USA, #634413), analyzed on Agilent TapeStation 4200 (Agilent Technologies, Inc., Santa Clara, CA, USA, #G2991AA), and pooled before paired-end 150-base-pair sequencing on one lane of an Illumina NovaSeq 6000 sequencer (Illumina, Inc., San Diego, USA), which generated 3 million reads per samples. Salmon (v0.13.1) was used to map reads to the human transcriptome (Gencode v28). Tximport (v1.14.0) was used to import the data into R (v3.6.1) and DESeq2 (v1.26.0) was then used to perform differential gene expression analysis. For predicting the target pathways, iPAGE (v1.0) with default settings was used [13], which uses the concept of mutual information to directly quantify the dependency between the expression data set and known pathways in MSigDB.

### Statistical analysis

Statistical analysis, t-test, linear regression, and odds ratios with a Fisher’s exact test were performed on GraphPad Prism 8. Mean and standard error were reported. P-values less than or equal to □ 0.05 were considered significantly different.

## RESULTS

### The freeze-thaw process decreases cell outgrowth from postmortem dura mater

To investigate the potential of postmortem dura mater as a viable source of cultured fibroblasts, we performed a systematic comparison of the ability to culture cells from two different sources, dermis and dura mater. Dermal cells were obtained from skin biopsies during life and dura mater was either processed shortly after autopsy (fresh) or from banked frozen dura mater (frozen).

For this comparison, we processed a total of 77 dermal biopsies for fibroblast cell banking [23]. These samples were obtained from subjects during life with a mean age of 53.6 ± 14.2 years and with more female subjects than male subjects, 62% and 38% respectively (Table 1). Most subjects were diagnosed with neurogenerative disease, either clinically or through neuropathological assessment (Supplemental Table S1). All processed dermal cell lines produced cell outgrowth, and cell lines were successfully expanded and banked in liquid nitrogen (Table 2). Thus, the generation of cell lines from dermal biopsies performed in living subjects was robust and consistent.

**Table 1.** Demographics of dermal and dural cases

|  | **Dermis** | **Dura mater** |  |
| --- | --- | --- | --- |
| **Mean** | **Fresh** | **Fresh** | **Frozen** |
| Number of cases | 77 | 43 | 14 |
| Age (years) | 53.6 ± 14.2 | 74 ± 9.3 | 66 ± 8.5 |
| PMI (hours) | - | 11.1 ± 7.6 | 7.4 ± 2.2 |
| Sex (M/F) | 29/48 | 23/20 | 7/7 |

**Table 2.** Successful cell outgrowth and karyotype (KT) analysis of dermal and dural cases

|  | **Dermis** | **Dura mater** |  |
| --- | --- | --- | --- |
|  | **Fresh** | **Fresh** | **Frozen** |
| Outgrowth | 77/77 (100%) | 40/43 (93%) | 2/14 (14%) |
| Normal KT | 31/36 (86%) | 12/29 (41%) | 2/2 (100%) |
| Abnormal KT | 5/36 (14%) | 13/29 (45%) | 0/2 (0%) |
| Unstable KT | 0/36 (0%) | 4/29 (14%) | 0/2 (0%) |

To compare the success rate of cell outgrowth from dura mater, we utilized the same isolation protocol as for dermal cell banking. We processed a total of 43 fresh dural cases, from subjects with a mean age of 74 ± 9.3 years, mean postmortem interval (PMI) of 11.1 ± 7.6 hours, and with a slightly greater number of male subjects than female subjects (53% and 47% respectively, Table 1). Most subjects were diagnosed with neurogenerative disease through neuropathological assessment (Supplemental Table S1 and Table S2). Forty cases had cell outgrowth that were successfully banked (93%, Table 2). Furthermore, two dural cases that initially had no cell outgrowth were recovered by reprocessing tissue that was cryoprotected with DMSO and stored in liquid nitrogen post-autopsy. Three fresh dural cases displayed no outgrowth, despite extensive time in culture (∼ 40 days). In summary, cell outgrowth, generation, and banking from fresh dural tissue were comparable to the dermal tissue.

In addition, we investigated the ability to culture cells from banked frozen dura mater. A total of 14 frozen cases were processed, with the mean age of subjects being 66 ± 8.5 years, mean PMI of 7.4 ± 2.2 hours, and male and female subjects were equally represented (Table 1). Only two frozen dural cases had cell growth that was sufficient for successful banking (14%, Table 2). Three additional dural cases had limited cell outgrowth, slow proliferation, and poor cell morphology and therefore could not be successfully banked. In an attempt to increase cell recovery from frozen dura mater, we utilized several alternative culture media for seven frozen dural cases [3, 22]. Despite up to 40 to 70 days in culture, cell outgrowth from the different types of media compositions and combinations could not be consistently improved. Overall, the fresh dural tissue showed significantly better cell outgrowth compared to frozen dural tissue (with an odds ratio of 80, 95% CI [10.71-407.9], p< 0.0001****, Fisher’s exact test).

### Dura mater-derived cell lines exhibit slower outgrowth and proliferation, as well as abnormal morphology

As dermis and dura mater-derived cell lines were cultured and banked, differences in initial cell outgrowth from the tissue, cell morphology, and proliferation rates became apparent. From dermal tissue, cell outgrowth was observed by day seven post-dissection and an adequate number of cells were expanded for cryopreservation after thirty days. In contrast, cell outgrowth from fresh dura mater was observed approximately nine days post-dissection and cells were cryopreserved after thirty-one days. Finally, cells derived from frozen dural tissue took 19 days for outgrowth post-dissection and were cryopreserved after 55 days (Fig. 1). Thus, initial cell outgrowth and banking from dural tissue took longer compared to dermal tissue.

**Figure 1.**
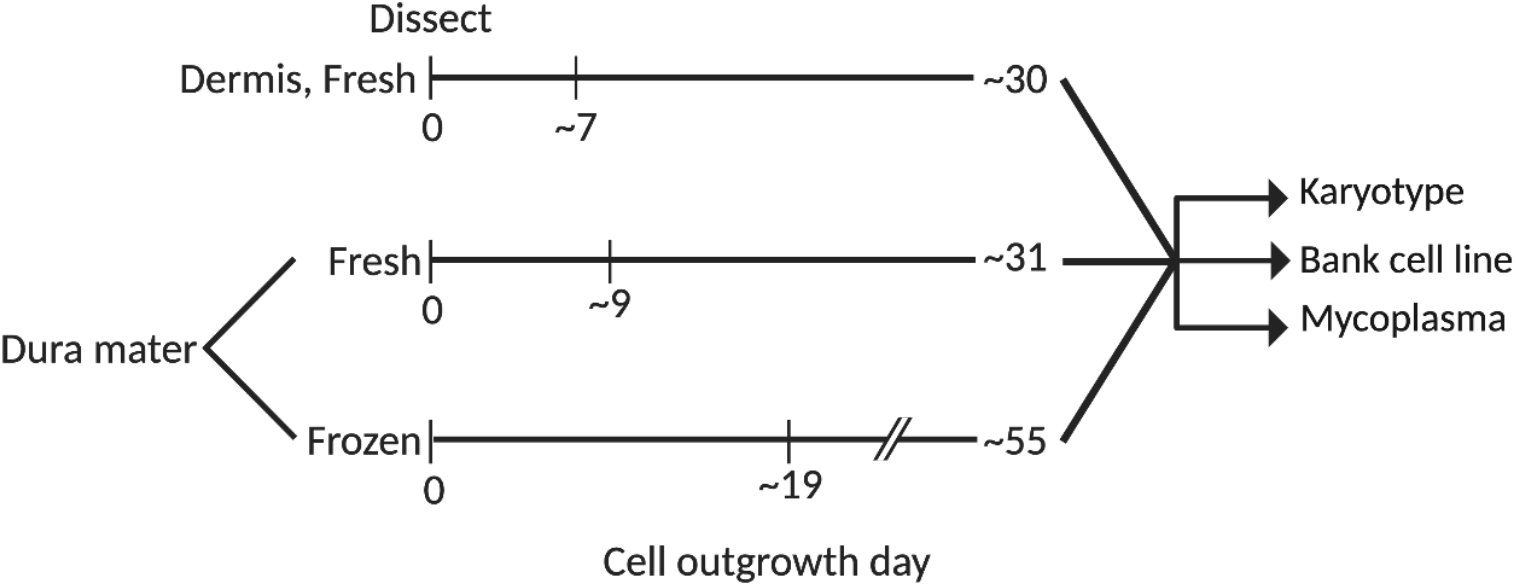
Timeline in days for the isolation, expansion, and banking of dermal cells from fresh dermal and dural cells from fresh and frozen dura mater. Cell outgrowth, proliferation, and banking from fresh and frozen dural tissue were delayed compared to dermal tissue. Successfully banked cell lines were karyotyped and tested for mycoplasma.

Differences in cell morphology between the dermis and dura mater-derived cell lines were also noted. Brightfield images demonstrated that dermis-derived cells exhibited classical fibroblast morphology, with long, spiny, and tightly packed cell bodies. Dural cells derived from fresh dura mater appeared to have an intermediate phenotype, with some normally spindle-shaped cell bodies that were densely packed and others with a more cobblestone shape and disorganized alignment. In contrast, dural cells derived from frozen dura mater displayed abnormal morphological features, such as enlarged cell bodies, decreased uniformity, and decreased cell density (Fig. 2A). Rhodamine phalloidin staining for actin filaments further illustrated the decreased organization of dura mater-derived cell lines (Fig. 2B). Taken together, these observations show differences in cell morphology and arrangement between dermis and dura mater-derived cells, suggesting potential abnormalities in the dura mater-derived cell lines.

**Figure 2.**
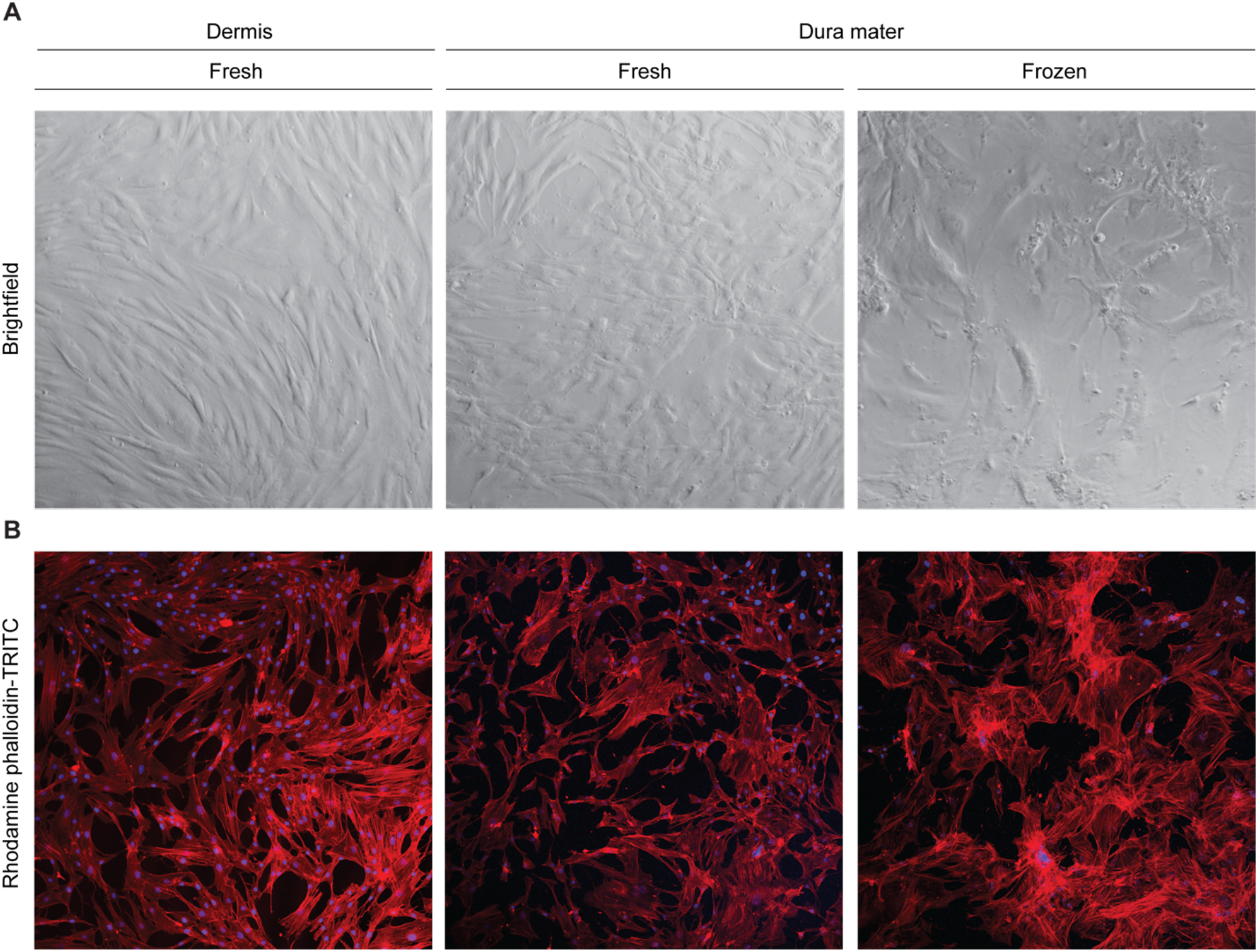
Dural cells displayed reduced cell outgrowth with larger cell body morphology than cultured dermal cells. **A** Dermal cells exhibit classical fibroblast morphology, with long, spiny, and tightly packed cell bodies. In contrast, dural cells are enlarged, variable, and have disorganized alignment. Dermal and frozen dural cells are from the same subject. Representative brightfield images are shown for each cell type and condition with a 5x objective **B** Cells were stained with rhodamine phalloidin conjugated to TRITC, reflecting differences in actin filament arrangement between dermal and dural cells. Representative immunofluorescent images are shown for each cell type and condition with a 10x objective.

Next, we documented proliferation rates of several lines. Compared to dura mater-derived cell lines, dermis-derived cell lines resulted in more total cells counted over the span of six days (Fig. 3A). Proliferation rate of the cell counts showed a significant decrease in dura mater-derived cell lines compared to dermis-derived cell lines (Fig. 3B). Overall, dermis-derived cell lines were observed to proliferate about three times faster than dura mater-derived cell lines.

**Figure 3.**
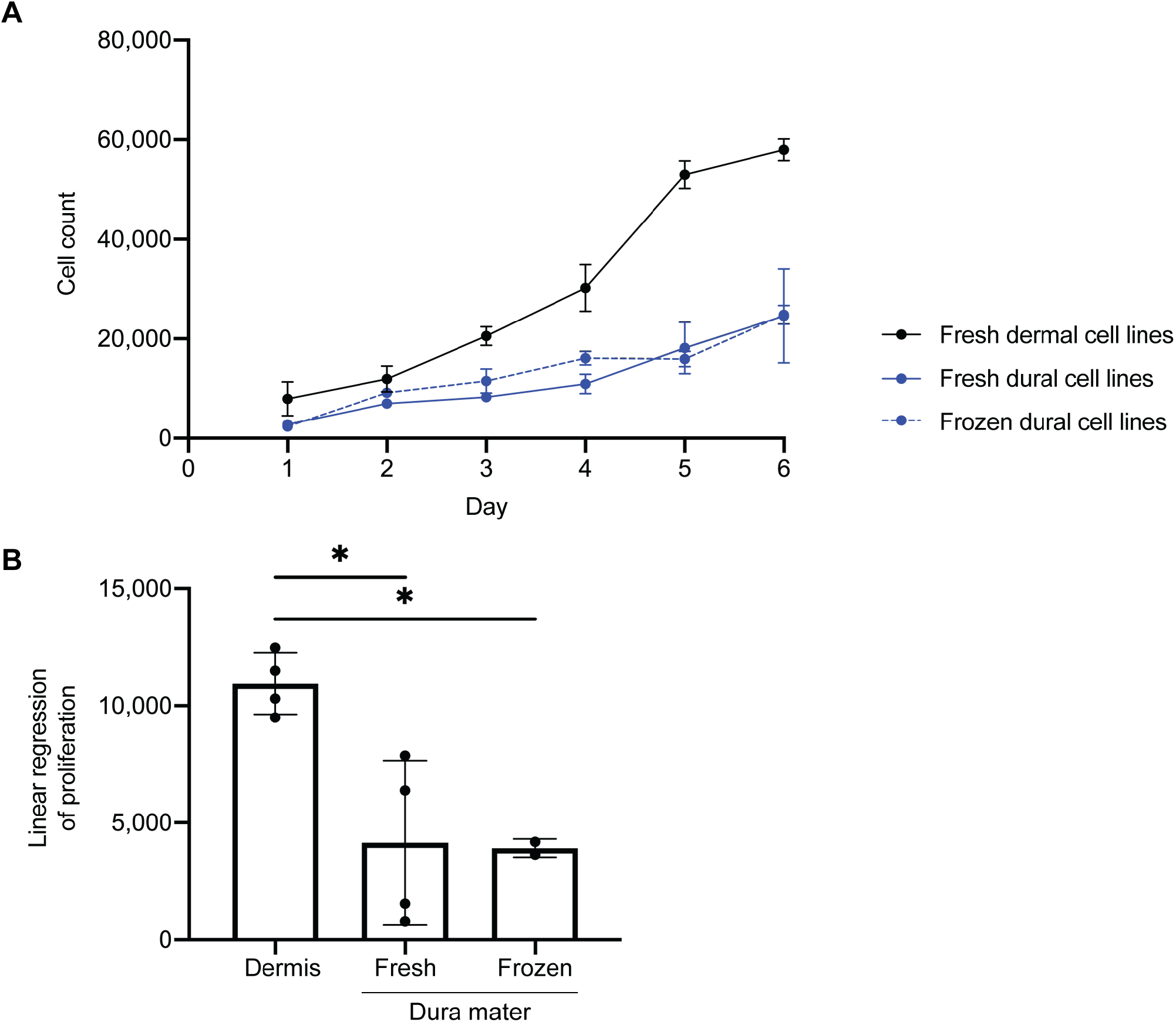
Differences in cell proliferation were observed between cells derived from dermis and dura mater. **A** Four dermal cell lines (black), four fresh dural cell lines (solid blue), and two frozen dural cell lines (dotted blue) were plated on day 0 and the total number of cells were counted every day for a span of six days. Total cell counts for each day are shown, performed in triplicates. **B** Cell proliferation rates for dermal and dural cell lines are shown. There was a significant decrease in cell proliferation in both frozen and fresh dural cell lines compared to the dermal cell lines (One-way ANOVA, * p < 0.05).

### Dura mater-derived cell lines exhibit chromosomal abnormalities

As an important quality control in cell banking, we next performed chromosomal karyotyping to assess gross genetic changes or anomalies in the derived cell lines. From the thirty-six dermis-derived cell lines that underwent karyotype analysis, thirty-one cell lines exhibited a normal karyotype (86%) and five cell lines exhibited an abnormal karyotype (14%). In contrast, from the twenty-nine fresh dura mater-derived cell lines that underwent karyotype analysis, twelve cell lines exhibited a normal karyotype (41%), thirteen cell lines exhibited an abnormal karyotype (45%), and four cell lines exhibited an unstable karyotype (14%). Of note, the two cell lines derived from frozen dura mater that were successfully banked both exhibited normal karyotypes (Table 2). Overall, dura mater-derived cell lines revealed significantly more chromosomal instability than dermis-derived lines.

We wondered whether sex of the donor affected karyotype. Interestingly, from the 13 dura mater-derived cell lines that exhibited abnormal karyotypes, 12 cell lines were derived from male subjects (92%). Furthermore, the majority of male-derived lines with chromosomal abnormalities exhibited the clonal loss of chromosome Y (LOY). There were eight cell lines that exhibited only LOY, three cell lines that exhibited LOY with additional autosomal abnormalities, and one cell line that exhibited trisomy in chromosome 7 (Table 3). In summary, there were more chromosomal abnormalities in male-derived dural cell lines compared to female-derived dural cell lines (with an odds ratio of 16.8, 95% CI [1.615-201.2], p = 0.0112*, Fisher’s exact test).

**Table 3.** Chromosomal abnormalities of dural cell lines from fresh dura mater

|  | **Male (n = 12)** | **Female (n = 1)** |
| --- | --- | --- |
| Trisomy | 1 | 1 |
| LOY | 8 | - |
| LOY with autosomal abnormalities | 3 | - |

LOY = Loss of chromosome Y

### Dura mater-derived cell lines fail to express key fibroblast markers

Given the differences in morphology, proliferation rate, and karyotype between dermal and dural cells, we next asked questions about the cell identity of dermal versus dural cell lines (Supplementary Table S3). We assessed cell identity by immunostaining against two classical markers of fibroblast identity, the intermediate filament protein vimentin and the fibroblast-specific protein 1 (FSP1 or S100A4), a member of the S100 calcium binding family [16] (Fig. 4A). While dura mater-derived cells expressed vimentin, albeit at lower levels, they strikingly lacked expression of S100A4 (Fig. 4A-C). In this analysis, we were able to compare protein expression in two pairs of dermis and dura mater-derived cells from the same subjects (Figure 4A, lanes 1A/1B and 2A/2B). Notably, in these “matched” cell lines, levels of the S100A4 fibroblast marker appeared decreased in the cells derived from postmortem dura mater compared to cells derived from dermis. Therefore, two fibroblast markers differed even between dermis and dura mater-derived cell lines, even when the cell lines were generated from the same subject.

**Figure 4.**
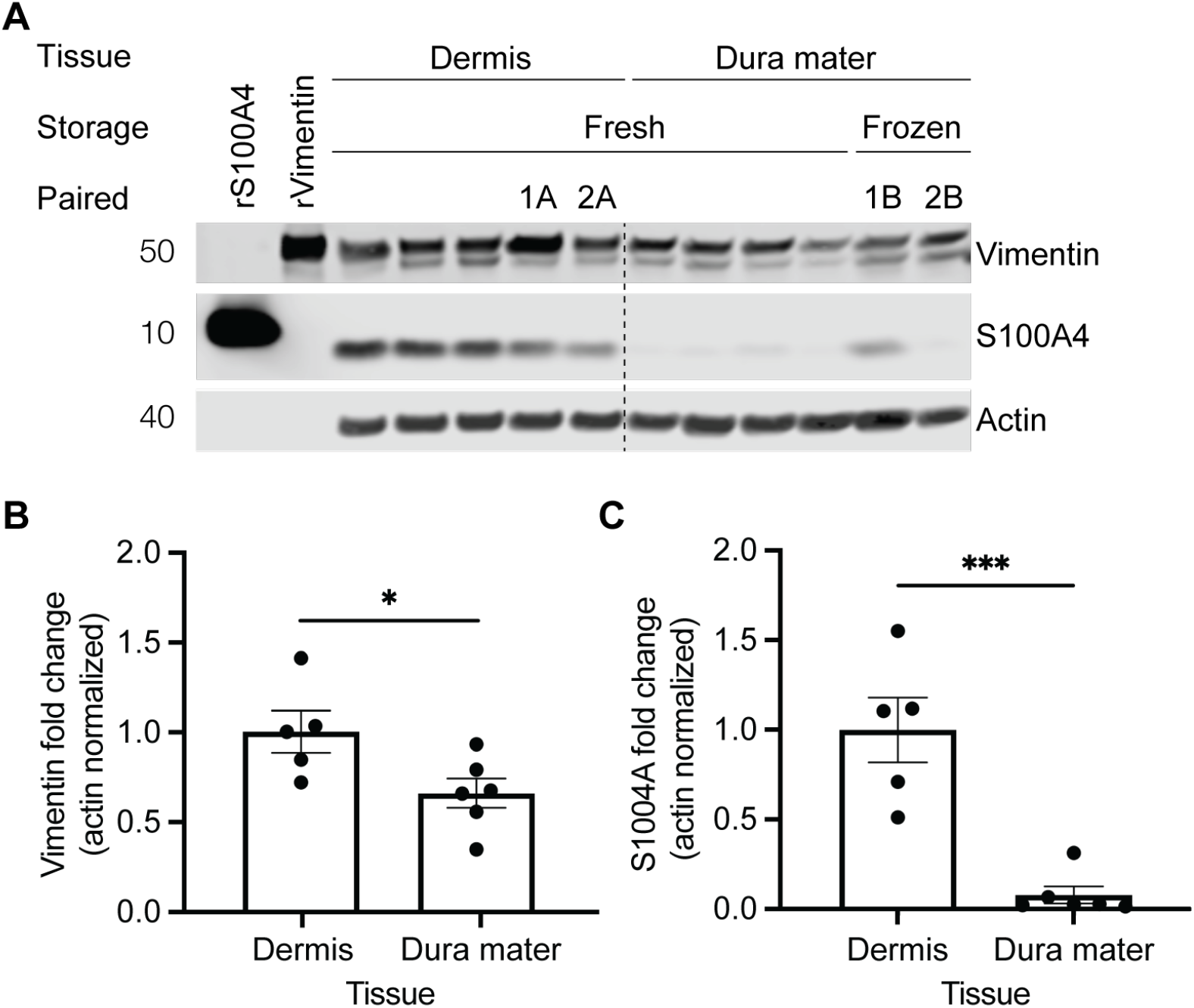
Protein expression differences in fibroblast markers were observed between dermal and dural cell lines. **A** Cell lysates from five dermal and six dural cell lines (20 µg) were immunoblotted with anti-Vimentin and anti-S100A4 antibodies. Tissue indicates from which source the cells were derived from, storage indicates whether tissue underwent frozen or fresh storage conditions before processing, and paired indicates cell lines derived from the same subject (2 cases, lanes 1A/1B and 2A/2B). Actin was used as a loading control. rVimentin and rS100A4 were recombinant proteins that were used as positive controls for their respective antibodies. **B** There was a significant decrease in vimentin protein levels in dural cell lines relative to the dermal cell lines (normalized to actin, unpaired t-test, * p < 0.05). **C** There was a significant decrease in S100A4 protein levels in dural cell lines relative to the dermal cell lines (normalized to actin, unpaired t-test, *** p < 0.0005).

### Dermis and dura mater-derived cells from the same subject exhibit highly divergent gene expression profiles

To better understand how cells cultured from dermal and dural tissues compare in an unbiased manner, we investigated their transcriptional profiles by RNA-sequencing. For this analysis, we utilized dermis and dura mater-derived cells from the same subject (lanes 1A and 1B from Fig. 4A). Notably, by definition, these cells were isolated at two different points in time. The dermal biopsy occurred during life when the subject was age 48 and the dura mater was frozen at the time of autopsy at age 51. RNA-sequencing analysis revealed approximately 3,000 genes that were significantly differentially expressed between the dermis and dura mater-derived cell lines. Gene ontology-term enrichment analysis revealed categories of genes significantly upregulated in dura mater-derived cells involved in cell adhesion, cell migration and proliferation, extracellular matrix and structure, filaments, and signaling. Genes associated with fibroblast identity that were upregulated in dura mater-derived cells included *ACTA2, ITGA1*, and *P4HA1*. Significantly downregulated genes were also found in the same categories: cell adhesion, cell migration and proliferation, extracellular matrix and structure, and filaments. Fibroblast associated genes that were downregulated in dura mater-derived cells included *FAP* and *ANPEP* (Fig. 5A, Additional File 1).

**Figure 5.**
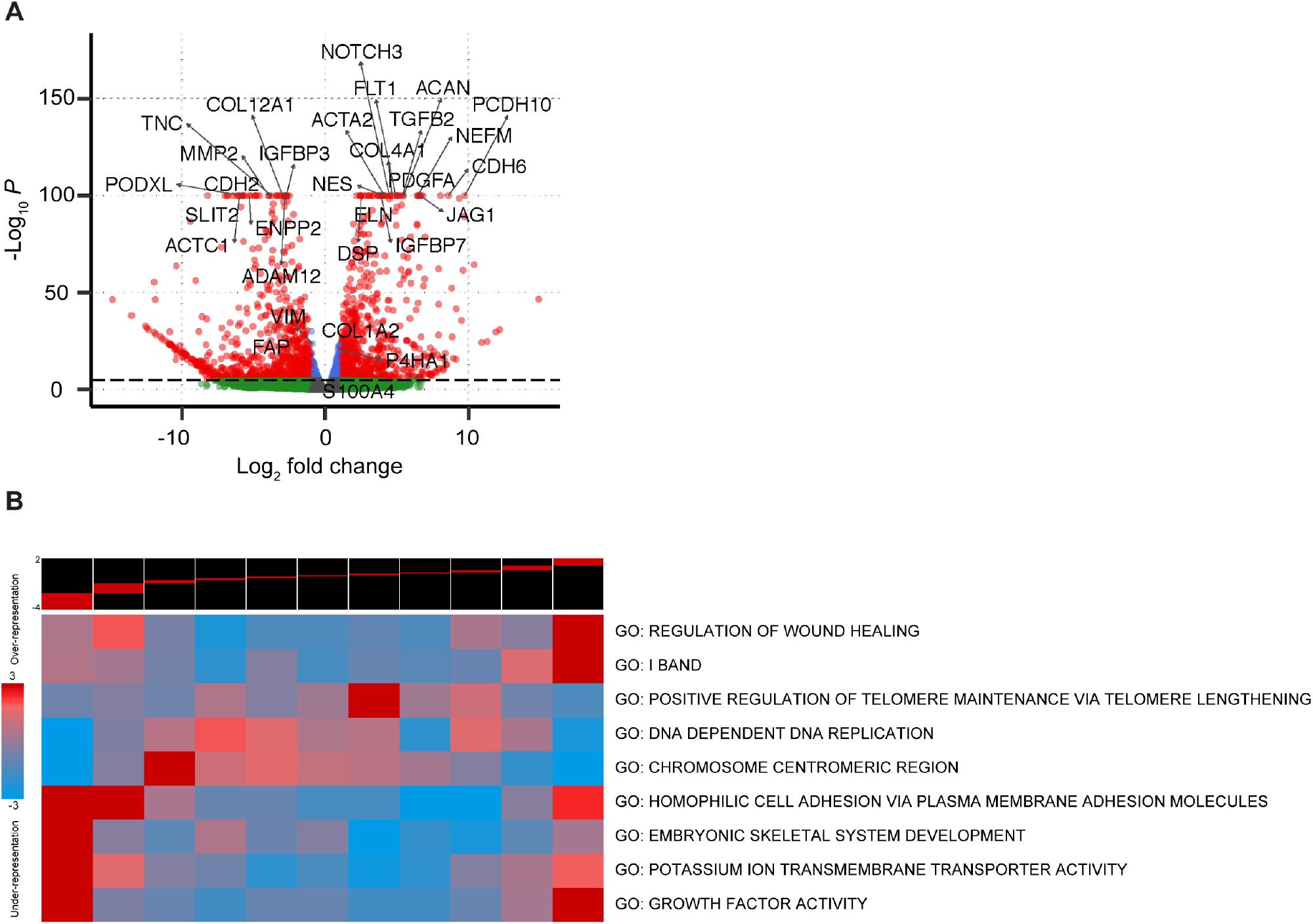
RNA-sequencing analysis of differentially expressed genes in dermal versus dural cells from the same subject. **A** Volcano plot of differentially expressed genes. Red dots denote differentially expressed genes, blue dots denote genes with expression differences of -Log_10_*P* between -1 and 1, and the green dots denote non-significantly expressed genes. **B** iPAGE pathways representation of differentially expressed genes. Top panel shows expression bins of genes with lower to higher expression in the dural cell line. Heatmap below indicates expression levels relative to GO terms. Red indicates enrichment/over-representation of pathway genes and blue indicates depletion/under-representation.

Given the number of differentially expressed genes, we further investigated implicated molecular pathways between the dermis and dura mater-derived cell lines through iPAGE and gene-set enrichment analyses. This analysis revealed upregulated pathways in categories such as “Regulation of wound healing”, which included cell shape, “I-band” which included actin filament, binding, and organization, and “Embryonic skeletal system development” which included cell fate specification, axis elongation, and mesenchymal to epithelial transition. iPAGE analysis also showed dysregulation in “Homophilic cell adhesions via plasma membrane” which included cell to cell adhesion and contact zone, “Potassium ion transmembrane transporter” which included actin filament movement, and “Growth factor activity” which included cell proliferation (Fig. 5B). Together, these data suggest that the dura mater-derived cells significantly differ from dermis-derived fibroblasts in both protein and gene expression, despite being derived from the same subject.

## Discussion

Fibroblast-like cells isolated from dura mater have been generated and reprogrammed into iPSCs and differentiated into neurons [2,3,22]. Postmortem dura mater-derived cells are advantageous when no other cell source is available. With the hope of advancing dura mater-derived cells as a potential cell source for precision medicine applications, we undertook a systematic comparison between cells isolated from dermal biopsies during life versus postmortem dura mater, either fresh from autopsy or after a period of being frozen. Earlier studies have characterized dura mater cells either collected perioperatively [3] or from postmortem tissue [2, 22]. In the latter cases, the dura mater cells were compared to cells derived from postmortem scalp dermis [2,22]. In contrast, this study compared postmortem dural cells to dermal fibroblasts collected during life. A comparison of the study materials, conditions used, and results reported in this and the prior dura mater cell studies can be found in ‘Additional file 2’. Our results revealed substantive differences between dermis and dura mater-derived cell lines in cell outgrowth, proliferation, morphology, karyotype, protein expression and transcriptional profiles. Taken together, these results raise questions regarding the “fibroblast” identity of dura mater-derived cells.

Some of the differences between our results and those of other groups may be due to technical differences. For example, Sproul et al. reported a higher success rate of fibroblast outgrowth from dura mater, which was processed by freezing in a liquid nitrogen vapor sandwich, with the tissue first chilled between aluminum plates and then placed in liquid nitrogen vapor for five minutes [22]. In comparison, we froze dura mater using a dry ice sandwich method, which is significantly warmer than liquid nitrogen vapor [24]. With these freezing conditions and the lack of a cryoprotectant reagent to prevent ice damage, the frozen dura mater and dural cells may have been damaged, resulting in the unsuccessful establishment of working cell lines or altered morphology. Therefore, careful storage precautions should be taken into consideration while freezing dura mater without a cryoprotectant reagent.

Cell outgrowth issues were ameliorated in dura mater that was processed fresh, without being frozen. Despite this, cell lines from fresh dura mater still contained a higher percentage of chromosomal abnormalities when compared to dermal fibroblasts. This observation could stem from several possibilities, including postmortem tissue source, age of subject, PMI, or more advanced stage of neurodegenerative disease. A significant proportion of the chromosomal abnormalities in dura mater-derived cell lines were from male cases that exhibited LOY. Interestingly, it has been previously reported that LOY in blood cells was associated in aging men with diseases, such as cancer and Alzheimer’s disease [8]. This may suggest that postmortem dura mater-derived cell lines from subjects with neurodegenerative diseases have chromosomal instability *in vitro*.

Sproul et al. also previously reported LOY in a dura mater-derived iPSC line which they attributed to *in vitro* culturing because the Y chromosome was present in the original frozen dura mater through *AMG* gene amplification [22]. This may be the case here as well. Although, another study generated a cell line from dura mater obtained during surgery and generated an iPSC line with a normal male karyotype [3]. Furthermore, it has been previously reported that derived human stem cells with chromosomal abnormalities have a cellular growth advantage *in vitro*, which may be similar within the dura mater-derived lines. The mechanisms of chromosomal abnormalities are unclear [1].

While karyotype abnormalities appear to be a significant concern for dura mater-derived cells, dural cell lines with a normal karyotype nonetheless did not exhibit a clear fibroblast identity based on protein markers and RNA-sequencing profile expression. Expression differences in cell lines derived from dermis and dura mater may be due to the different tissue types and anatomical locations as has been reported for fibroblasts isolated from different body regions such as the arm or thigh [4,7]. Others have reported a lack or decrease of S100 family protein expression in dura mater-derived cells [5,21,22], similar to what we have shown here. Thus, our data suggest that cells cultured from dura mater may not be fibroblasts but rather an alternative, yet unidentified cell type. As such, use of dermal cells for disease modeling should proceed with caution.

## Conclusions

We aimed to assess the possibility of utilizing postmortem dura mater, from both frozen and fresh storage conditions, as a source of *in vitro* disease cell models. Contrary to our expectations, this study revealed multiple phenotypic, karyotypic, and transcriptional differences between dermis and dura mater-derived cells, even when those cells originated from the same subject. The reasons underlying these differential features are not entirely clear. However, this study should prompt, at a minimum, careful karyotyping and functional assessment of dura mater-derived cells prior to reprogramming into iPSCs. Additionally, further investigation into the overall utility of dura mater-derived cells as a source for disease modeling is warranted.

## Additional Files

## List of abbreviations

PMI: postmortem interval
iPSCs: induced pluripotent stem cells
DPBS: Dulbecco’s phosphate buffered saline
KT: karyotype
LOY: loss of chromosome Y
S100A4/FSP1: fibroblast-specific protein 1
rVimentin: recombinant vimentin
rS100A4: recombinant S100A4

## Ethics approval and consent to participate

Consent for brain donation was obtained from all subjects or their surrogates in accordance with the Declaration of Helsinki and the research was approved by the UCSF Committee on Human Research. Dermal biopsies were obtained from subjects with written informed consent and the research was approved by the UCSF Committee on Human Research (#10-00234).

## Consent for publication

Not applicable.

## Availability of data and material

Cell lines are available upon reasonable request through the Memory and Aging Center at the University of California, San Francisco. A detailed dura mater-derived cell culture protocol can be found at protocols.io: dx.doi.org/10.17504/protocols.io.8m2hu8e. RNA-sequencing data from this study has been deposited to Gene Expression Omnibus.

## Authors’ contributions

ARA, WWS, and AWK planned and designed the study. ARA, KG, CIC, and HG performed the experiments. CGC, ALN and AMK provided tissue and supporting information. SS, LTG, and WWS performed neuropathological assessment. ARA, KG, and HG compiled and analyzed the data. ARA and AWK composed the manuscript. All authors read and approved the final manuscript.

## Acknowledgements

The authors thank the members of the Kao and Seeley lab for their helpful discussions and support in the laboratory. The authors thank the members of the CZI Neurodegeneration Challenge Network for their support and feedback. The authors gratefully acknowledge and thank the funding that supported this study. We thank the participants and their families for their contributions to this study and research.

---

**Table S1.** Diagnosis of dermal and dural cases

|  | **Dermis** | **Dura mater** |  |
| --- | --- | --- | --- |
|  | **Fresh (n = 77)** | **Fresh (n = 43)** | **Frozen (n = 14)** |
| Neuropathological diagnoses of disease | 8 (10%) | 38 (88%) | 14 (100%) |
| Clinical diagnoses of disease | 51 (66%) | - | - |
| Clinically normal | 14 (18%) | 5 (12%) | - |
| Biospecimen only | 4 (5%) | - | - |

**Table S2.**
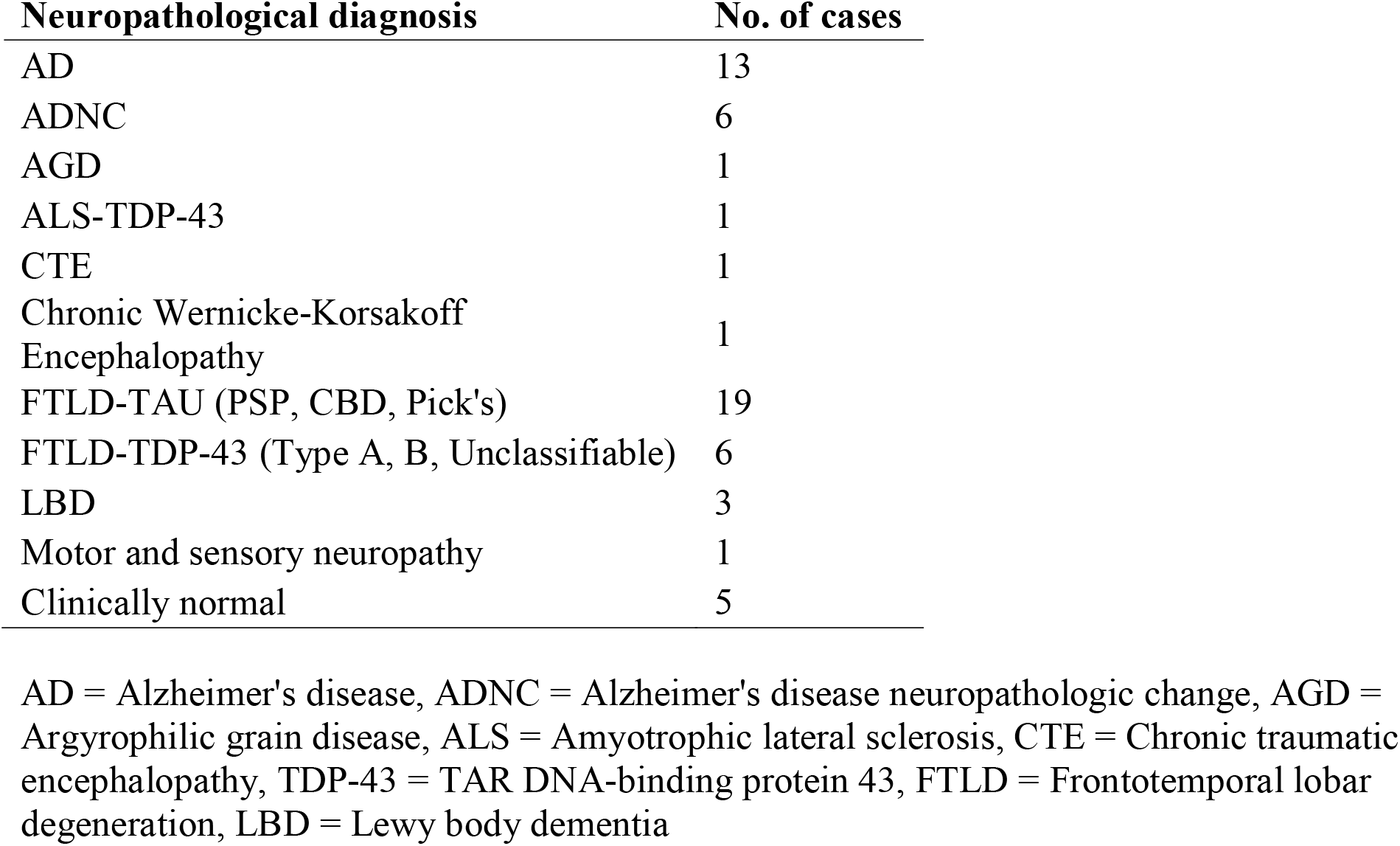
Neuropathological diagnosis of all dural cases

| **Neuropathological diagnosis** | **No. of cases** |
| --- | --- |
| AD | 13 |
| ADNC | 6 |
| AGD | 1 |
| ALS-TDP-43 | 1 |
| CTE | 1 |
| Chronic Wernicke-Korsakoff<br>Encephalopathy | 1 |
| FTLD-TAU (PSP, CBD, Pick's) | 19 |
| FTLD-TDP-43 (Type A, B, Unclassifiable) | 6 |
| LBD | 3 |
| Motor and sensory neuropathy | 1 |
| Clinically normal | 5 |

AD = Alzheimer's disease, ADNC = Alzheimer's disease neuropathologic change, AGD = Argyrophilic grain disease, ALS = Amyotrophic lateral sclerosis, CTE = Chronic traumatic encephalopathy, TDP-43 = TAR DNA-binding protein 43, FTLD = Frontotemporal lobar degeneration, LBD = Lewy body dementia

**Table S3.** Selected dermal and dural cell lines for western blot analysis

| **Lane** | **Derived tissue** | **Tissue arrival** | **Age at biopsy** | **Age at death** | **PMI** | **Sex** |
| --- | --- | --- | --- | --- | --- | --- |
| 1 | Dermis | Fresh | 72 | - | - | M |
| 2 | Dermis | Fresh | 52 | - | - | F |
| 3 | Dermis | Fresh | 67 | - | - | F |
| 4<sup>1A</sup> | Dermis | Fresh | 48 | - | - | F |
| 5<sup>2A</sup> | Dermis | Fresh | 65 | - | - | F |
| 6 | Dura mater | Fresh | - | 72 | 28.2 | M |
| 7 | Dura mater | Fresh | - | 69 | 11.6 | M |
| 8 | Dura mater | Fresh | - | 60 | 20.5 | F |
| 9 | Dura mater | Fresh | - | 82 | 9.6 | F |
| 10<sup>1B</sup> | Dura mater | Frozen | - | 51 | 5.4 | F |
| 11<sup>2B</sup> | Dura mater | Frozen | - | 66 | 7.7 | F |

Dermal cell line 1A and dural cell line 1B are from the same subject.
Dermal cell line 2A and dural cell line 2B are from the same subject.

